# Sediment Selection: Range Expanding Fiddler Crabs are Better Burrowers Than Their Historic Range Counterparts

**DOI:** 10.1101/2020.11.14.351007

**Authors:** Richard J Wong, Michael S. Roy, Jarrett E. K. Byrnes

## Abstract

Species ranges are shifting in response to climate change. In New England saltmarshes, the mud fiddler crab, *Minuca pugnax*, is expanding north of Cape Cod, MA into the Gulf of Maine (GoM) due to warming waters. The burrowing lifestyle of *M. pugnax* means sediment compaction in saltmarshes may influence the ability for crabs to dig, with more compact soils being resilient to burrowing. Previous studies indicate that saltmarshes along the GoM have higher soil strength (i.e., compaction) relative to marshes south of Cape Cod. Together, physical characteristics and temperature of this habitat may be influencing the burrowing performance of *M. pugnax*, and therefore the continuation of their northward range expansion into the GoM. To determine if compaction affects burrowing activity of *M. pugnax* in historical and range expanded populations, we conducted a controlled laboratory experiment. We manipulated soil compaction in standardized lab assays and measured crab burrowing performance with individuals collected from Nantucket (i.e., historical range) and the Plum Island Estuary (PIE, i.e., expanded range). We determined compaction negatively affected burrowing ability in crabs from both sites; however, crabs from PIE burrowed in higher soil compactions than Nantucket crabs. In addition, PIE crabs were more likely to burrow overall. We conclude that site level differences in compaction are likely altering burrowing behavior in the crab’s expanded range territory by way of phenotypic plasticity or rapid evolution. Our study demonstrates that non-temperature physical habitat traits can be as important as temperature in influencing climate driven range expansions.

## Introduction

Global distributions of species are shifting because of climate change (Crozier, 2004; Dawson, 2010; Ling, 2008; Johnson, 2014; Sanford, 2006). Temperature is a known barrier that creates many species’ range borders (Burrows et al., 2014). As temperatures shift, barriers to species’ ranges can be breached, leading to range expansions (Parmesan, 2006). Such range expansions are occurring with increasing frequency across a variety of taxa and ecosystems (Krehenwinkel, 2013; Rochlin, 2013; Taulman, 2014). As they shift their range, species – particularly consumers and ecosystem engineers – can have a large effect in these novel areas. For example, warming waters enabled the southward range expansion of the sea urchin *Centrostephanus rodgersii* in the Tasman Sea (Ling, 2008; Ling et al., 2009; Ling and Johnson, 2012), leading to profound consequences, from the denuding of kelp forests and competition-driven declines in abalone populations (Strain et al., 2013). Temperature, however, is not always the only barrier to species’ ranges (Burrows et al, 2014). Unsuitable habitat, lack of prey, and abundance of predators, to name a few, can serve to slow or even stop range expansions. In contrast, plasticity and rapid evolution could counteract this mismatch. How species survive, thrive, and change when expanding their ranges due to opening thermal windows, despite other forms of mismatch, remains relatively unexplored.

Species that are stymied or prevented from experiencing range shifts, despite favorable temperature conditions, are often the stopped by non-climate related physical barriers (Alof et al., 2015; Edwards et al., 2013; Spence & Tingley, 2020). Water chemistry and stream and lake morphology, for example, slowed the expansion of several sport fish species in Canada (Alof et al., 2015). These fish were physically incapable of entering and persisting in some Canadian freshwater bodies where their thermal thresholds were met (Alof et al., 2015). Rusty crayfish, *Orconectes rusticus*, are also experiencing northward range expansion facilitated by warmer climates (Phillips et al., 2009). *O. rusticus* cannot persist in the Canadian Shield, however, due to lower dissolved calcium (Edwards et al., 2013). Any small populations that do encroach deplete much of the remaining calcium, further arresting their poleward expansion (Edwards et al., 2013). These examples underscore the necessity of comprehending the full suite of conditions that facilitate shifts in species distributions.

We see a potentially similar story in the mud fiddler crab, *Minuca (Uca) pugnax*, in the Gulf of Maine. *M. pugnax* are expanding their range north due to warming waters but could be slowed by sediment characteristics. Although larvae have been in the GoM for the entirety of this species’ life history, crabs fail to develop in saltmarshes north of Cape Cod, MA, its historic northward range barrier. In 2003, Sanford et al. (2006) found mature *M. pugnax* north of the Cape. The proposed mechanism is a warming Gulf of Maine (GoM), which warmed at a rate that is twice the global mean in the last 40 years (Pershing et al., 2015). Sanford et al. (2006) demonstrated that *M. pugnax* larvae are highly sensitive to temperature; crab larval survival decreases exponentially as temperature falls below an 18°C threshold. As pelagic planktonic larvae, *M. pugnax* regularly experienced temperatures below this threshold in the GoM for most of its life history.

Although individuals were found north of the Cape by 2003, no individuals made it north of Boston, MA, until discovery of a population in the Plum Island Estuary (PIE) in 2014 (Johnson, 2014). The likely mechanism was an exceptionally warm summer 2012, where sea surface temperatures regularly exceeded 18°C well into the season of fiddler crab dispersal. Since 2014, *M. pugnax* has continued to expand northward along the New England coastline. Individuals have been found as far north as Hampton, NH (Johnson, 2014) and southern Maine (D. S. Johnson, *unpublished*). In addition, Sanford et al. (2006) found good gene mixing and adult winter temperature tolerances by crabs found north of the Cape but south of Boston, suggesting limited genetic or thermal barriers to expansion now that they have established populations. Crab larvae from the range edge also grow faster than crabs from south of Cape Cod (Sanford, 2006). This tolerance along with the genetic variability of northern populations suggests that the crabs are adapting to overcome physical barriers.

As a burrowing species of crab, *M. pugnax* could be particularly sensitive to the characteristics of saltmarsh sediment in its expanded versus historical ranges. In particular, soil compaction or soil density of novel marshes would likely influence the capacity of crabs to burrow; i.e., more compact sediments are more difficult to burrow into. We see this in other burrowing crabs, such as *Helice tientsinensis* in China, which have higher burrow densities in softer and wetter sediments than harder and drier sediments that are more difficult to burrow into (Li et al., 2018). Fiddler crabs burrow to feed, avoid predation, and to mate (Bertness & Miller, 1984; Luk & Zajac, 2013); therefore an inability to burrow would severely impact the basic life history characteristics of *M. pugnax*. Roy et al. (*unpublished, in prep*) demonstrated that compaction in marshes in Nantucket (i.e., historical range) were significantly lower (historical range, 13.8+/-0.871psi) than PIE (expanded range, 30.2+/-1.53psi, p<0.0001). Vincent (2013) also reported soil strengths of more than 50 psi (3.51 kg/cm^2^) in marshes along the GoM. In addition, fiddler crab densities are low in PIE (~3 crabs m^−2^) relative to saltmarshes south of Cape Cod (~150 crabs m^−2^) (Martínez-Soto & Johnson, 2020). The mechanism driving this low density could be due to reduced propagule pressure in expanded ranges, or some other physical barrier to survival and growth (e.g., difficult sediment to burrow into), or both.

To determine the relationship between physical substrate and crab burrowing ability in historical versus range expanded populations, we conducted a controlled laboratory mesocosm experiment of fiddler crab burrowing behavior in varying degrees of sediment compaction. In particular, we asked the following questions: 1) Does soil strength drive the capacity of fiddler crabs to burrow and influence the depth of fiddler crab burrows; 2) Are there differences in burrowing capacity and burrow depth between historical (i.e., Nantucket) versus expanded (i.e., PIE) fiddler crabs; and 3) Does fiddler crab population density play a role in burrowing behavior? We hypothesized that higher soil strengths negatively affect a crab’s burrowing performance (measured by burrow frequency and burrow volume). In other words, compact soil should impede burrowing crabs. We expected to see that, for crabs of similar sizes, both populations would be equally impacted by soil strength, indicating a role in higher soil strengths slowing range expansion, and no clear effect of density.

## 1. Materials and Methods

To evaluate the impact of soil strength on fiddler crab burrowing behavior, we developed a controlled laboratory mesocosm experiment testing burrowing behavior of crabs from natal versus range expanded populations in standardized soils varying in levels of compaction. Each mesocosm consisted of saltwater saturated peat moss compressed with varying weights of sand (see Supplementary Methods Figure 1). We combined 10 gallons of dry organic peat moss with 4 gallons of saltwater with a salinity of 20% in a 66 L bin to create our peat/salt mixture. After thoroughly mixing the peat, we left the mixture to sit overnight to fully saturate before using it for the experiment. To acclimate the crabs to the saturated peat moss environment, crabs were housed in clear 66 L bins with separately made peat moss/saltwater mixture.

We tested the following sediment compaction levels: 0 psi (0 kg/cm^2^), 10 psi (0.7 kg/cm^2^), 20 psi (1.4 kg/cm^2^), and 25 psi (1.8 kg/cm^2^). These values represent the average range of soil strengths Roy et al. (*unpublished, in prep*) measured in saltmarshes along the coast of Massachusetts in 2017 and 2018. We reached the desired soil strength for each treatment using a formula we developed to find the approximate amount of sand used (see Supplementary Methods and Supplementary Figure 1). We then placed the crabs in with the compacted soil for the trial (see Supplementary Methods Figure 2).

We tested two different crab densities to determine the effect of population density on burrowing behavior: one crab and three crabs per experimental chamber. Crabs were collected from Carolton Creek in Rowley, MA (42.745462, −70.836981) for the Plum Island trials and Folger’s Marsh in Nantucket, MA (41.294653, −70.041979) for the Nantucket trials (see Supplementary Methods Figure 3), both at daytime low tides. Trials were conducted close to the location where they were collected, except for one Plum Island Estuary trial conducted at the University of Massachusetts-Boston. We conducted four replications for each density and soil strength treatment. Total n = 128, n = 64 from each location (PIE and Nantucket), and n = 16 per replicate with four soil strength treatments and two different crab densities (one and three crabs).

Before each trial, crabs were randomly selected, weighed, and sexed, and then were put separately into their experimental vessel for four hours. After removing the crab(s), we counted burrows in each mesocosm, measured soil strength, and took plaster casts of each burrow found (see Figure 1). We determined burrow volume using water displacement by placing the cast in a graduated cylinder. Crabs were returned where they were collected after the experiment was completed.

**Figure 1:**
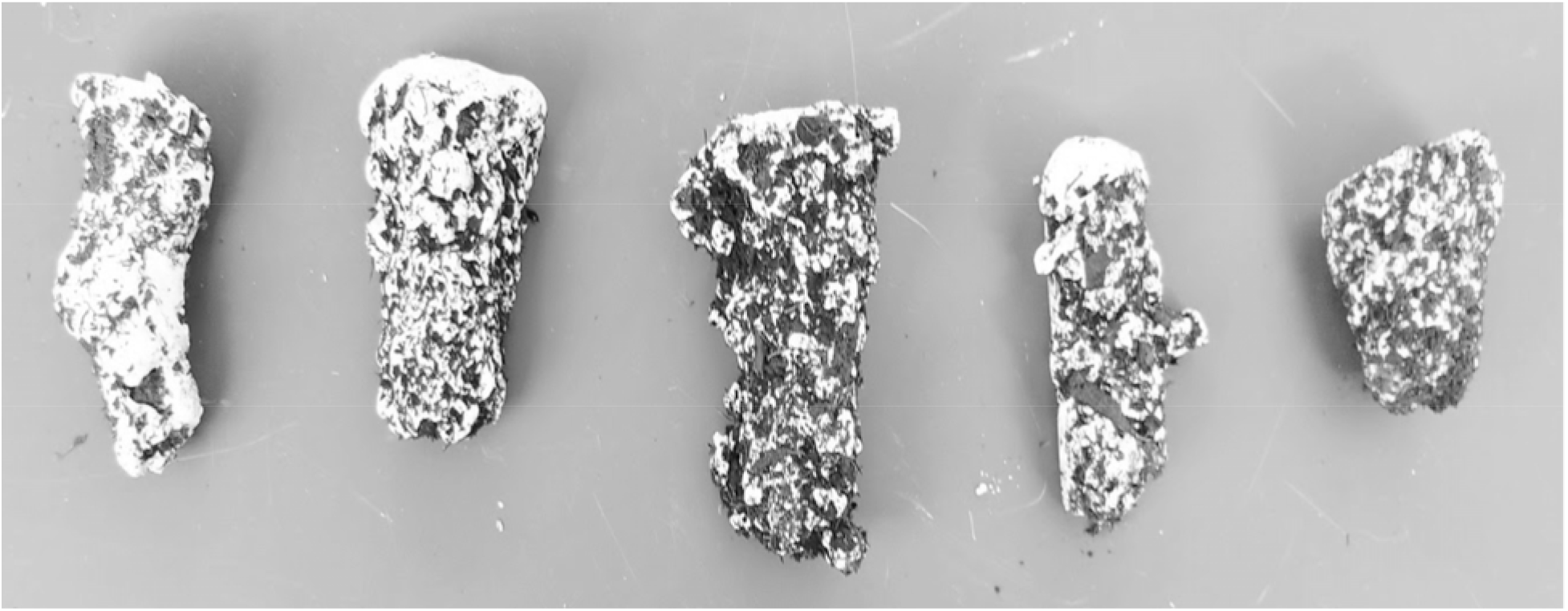
Example fiddler crab burrow casts. These were cleaned with a gentle toothbrush and placed into a graduated cylinder to find burrow volume.

To determine how soil strength influenced the probability of crab burrowing, we used binomial logistic regression (generalized linear model with logit link) with soil strength, crab natal location, their interaction, and crab mass as predictors and burrowing (yes/no) as a response. We fit separate models for the one and three crab experiments. To assess if soil strength affected burrow volume for those crabs that did burrow, we fit using a generalized linear model with a Gamma error and log link to accommodate for overdispersion and the lack of a 0 ml volume possibility. We used the same predictors and total burrow volume as a response. Note, using a gaussian error with an identity or log link produced the same results, but often led to impossible fitted values. All analyses were conducted in R version 3.6.1 (R Core Team 2019). All models were assessed for violations of assumptions using randomized quantile residuals using the DHARMa library (Florian Hartig, 2020). Code for all analyses can be found at https://github.com/richw1w/Pugnax_SS_Analysis.git

## 2. Results

Broadly, our results show that crabs collected in the Plum Island Estuary (expanded range) display better burrowing performance than Nantucket (historical range) crabs. In our one crab trials, Nantucket fiddler crabs were less likely to burrow at high soil strengths, whereas PIE fiddler crabs burrowed at all soil strengths (interaction effect, Figure 2A, Table 1, Supp. Table 1). Nantucket crabs did not dig at soil compaction levels past 10 psi, even though PIE crabs were able to burrow in all soil strengths (see Figure 3A). In three crab trials, crabs were less likely to burrow at higher soil strengths; PIE crabs in general have a higher probability of burrowing at all soil strengths (Table 1, Figure 2B) and, again, were the only crabs to burrow at >10 psi. Neither soil strength, site, their interaction, or any other predictor affected burrow volume of those crabs that did burrow (Table 2, Figure 3).

**Figure 2:**
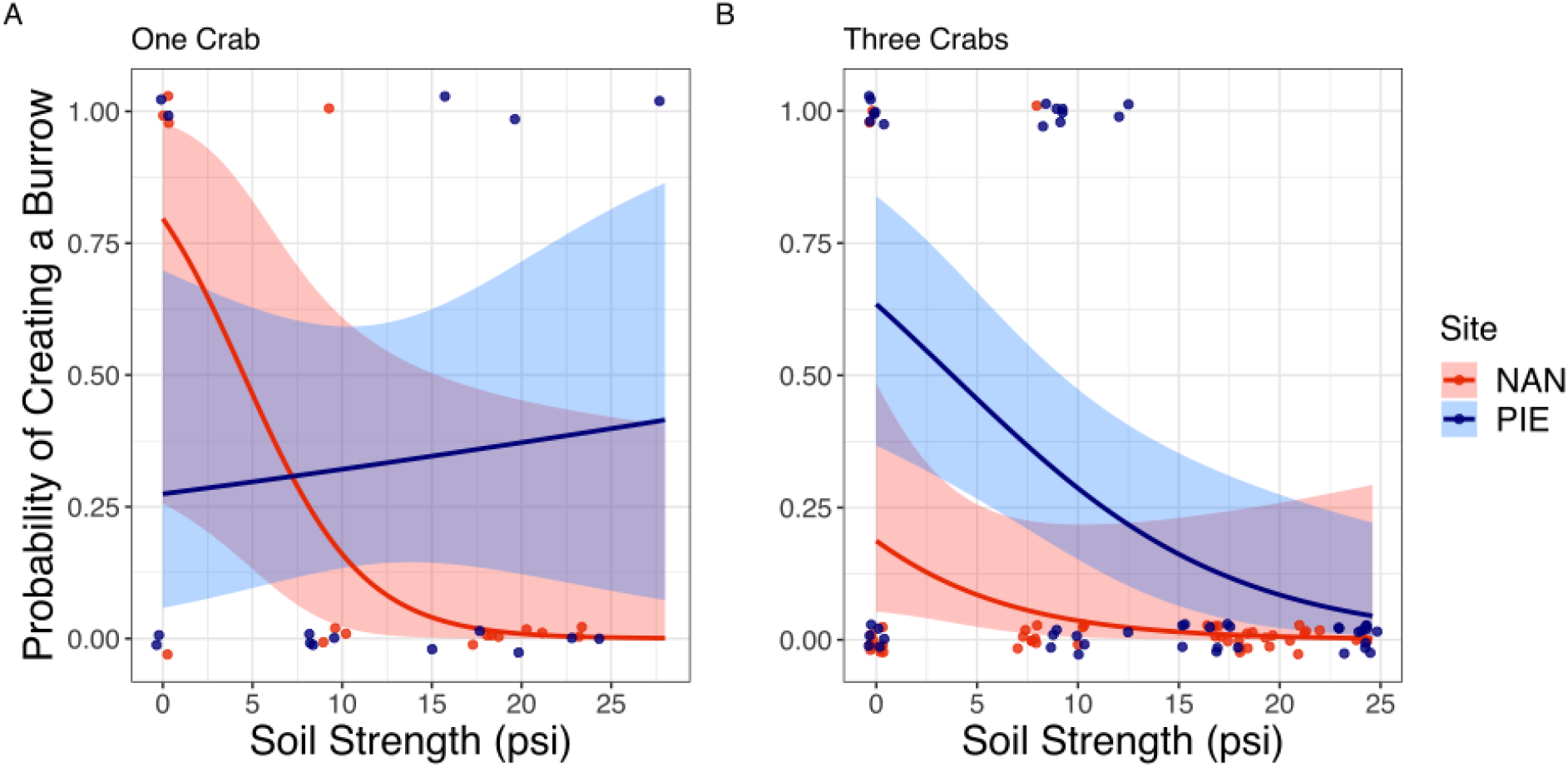
PIE crabs are more likely to burrow than those from Nantucket. The logistic relationship showing how soil strength affects the likelihood of the crab burrowing in trials with (A) one crab or (B) three crabs. Red represents Nantucket crab data, and blue represents PIE crab data. Curves are from fit models with 95% Confidence intervals. Points represent 1 = burrow or 0 = no burrow. Points are jittered in order to see overlapping data points and may not align exactly with the true data.

**Table 1:**
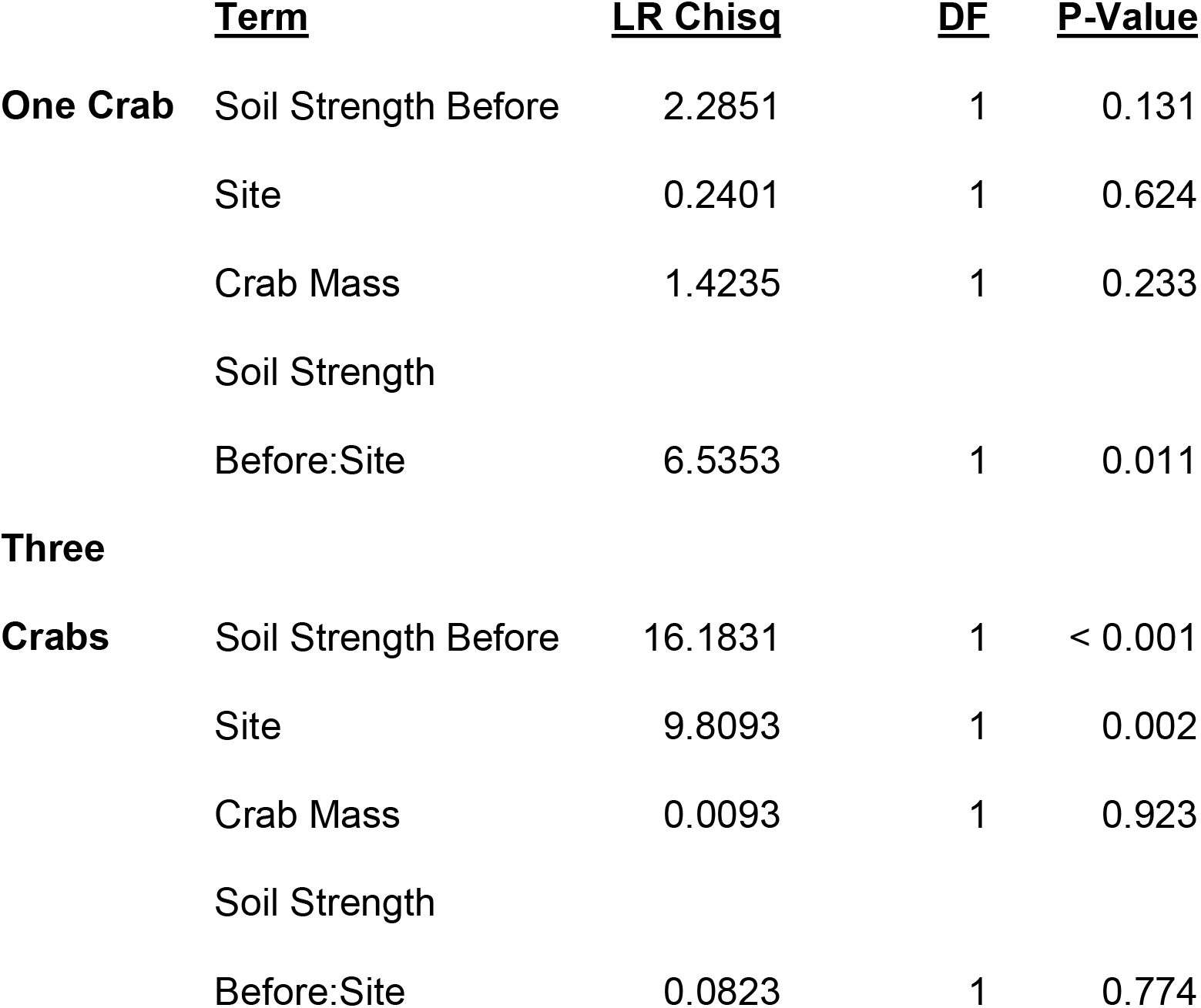
Analysis of Deviance results from probability of crab burrowing model for both the one crab trial and three crab trial.

**Figure 3:**
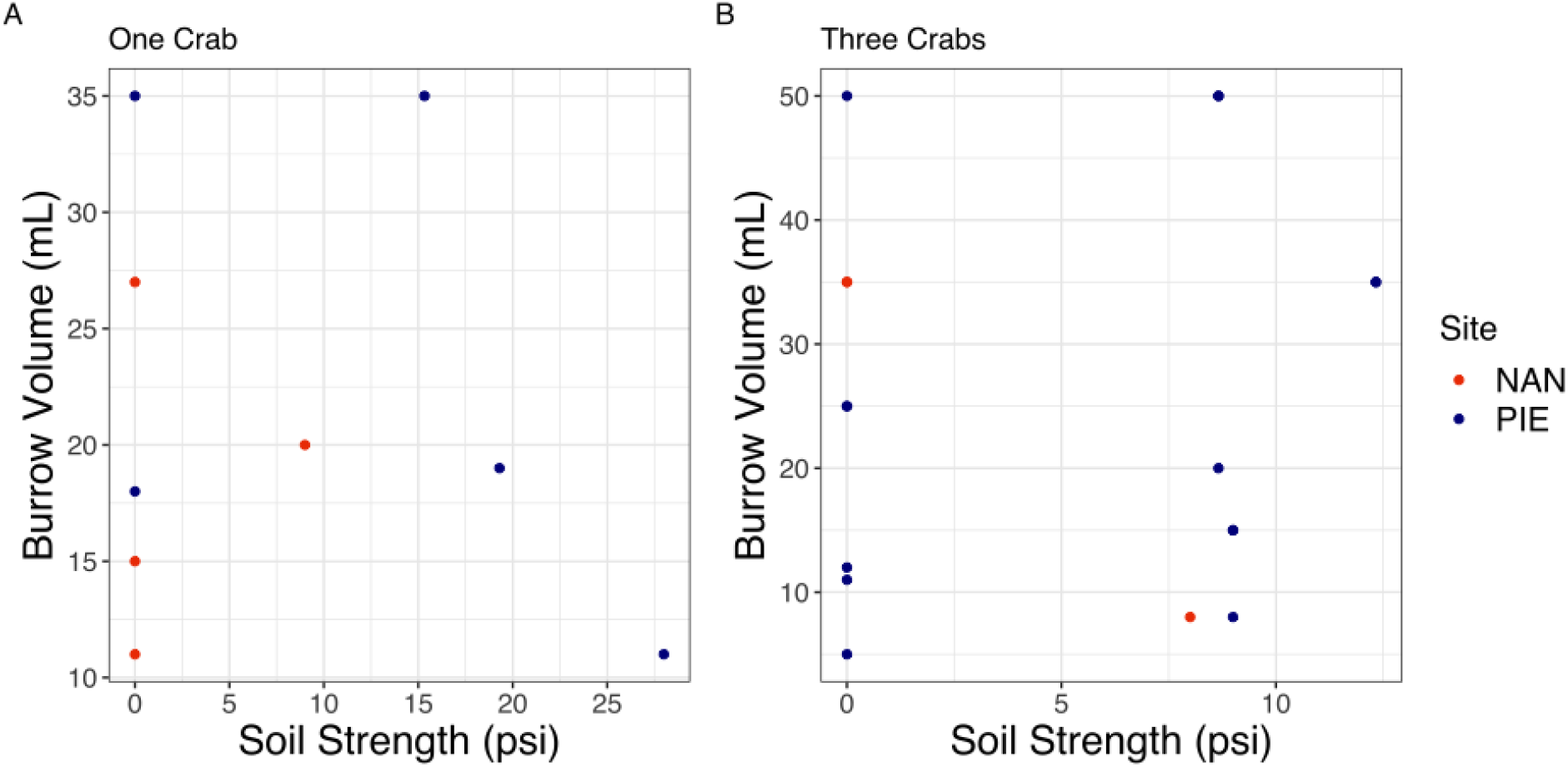
Nothing affects burrow volume in our experiment once a crab chooses to burrow. Data representing crab burrow volumes in trials with (A) one crab or (B) three crabs. Red represents Nantucket crab data, and blue represents PIE crab data. No curves are shown as no explanatory variables explained variability in the data.

**Table 2:** Analysis of Deviance results from probability of crab burrow volume model for both the one crab trial and three crab trial.

|  | <u>Term</u> | <u>LR Chisq</u> | <u>DF</u> | <u>P-Value</u> |
| --- | --- | --- | --- | --- |
| **One Crab** | Soil Strength Before | 0.9003 | 1 | 0.343 |
|  | Site | 1.0727 | 1 | 0.3 |
|  | Crab Mass | 0.1517 | 1 | 0.697 |
|  | Soil Strength |  |  |  |
|  | Before:Site | 0.3063 | 1 | 0.58 |
| **Three** |  |  |  |  |
| **Crabs** | Soil Strength Before | 0.0645 | 1 | 0.799 |
|  | Site | 0.089 | 1 | 0.766 |
|  | Crab Mass | 0.187 | 1 | 0.665 |
|  | Soil Strength |  |  |  |
|  | Before:Site | 2.9625 | 1 | 0.085 |

## 3. Discussion

In contrast to our initial expectations, our experiment shows that *M. pugnax* have likely changed their burrowing behaviors as they have expanded their ranges. We show that crabs living north of the Cape (e.g., PIE) are more capable of burrowing in compact soils than crabs living south of Cape Cod (e.g., Nantucket) (see Figure 2), less than 200 miles away (see Supplementary Methods Figure 3). In our trials with one crab per experimental chamber, PIE crabs were able to burrow in more compact soils than Nantucket (NAN) crabs (see Figure 3A). No Nantucket fiddler crabs were able to dig in soil strengths greater than 10 psi, however Plum Island Estuary crabs were able to dig in our densest soil treatment, 25 psi (see Figure 2A). These are directly comparable to the soil strengths that each population is experiencing in the environment from which they were collected (Roy et al.*, unpublished, in prep*). Even in trials with three crabs per chamber, PIE crabs were more likely to burrow than Nantucket crabs in all soil strength treatments (see Figure 3B). The difference between burrowing abilities in the two populations suggest that PIE crabs are stronger and able to dig in higher soil strengths. Although our results are from crabs originating in only one historical and one range expanded population, given soil strength differences south versus north of Cape Cod, we suggest this represents a more general trend. Our results do not speak to whether this is inherent genetic variation in the two populations driven by selection for stronger crabs or phenotypic plasticity of crabs settling in marshes with harder soils. Regardless, our results show that processes other than temperature have influenced this species as it has expanded its range.

Further, we suggest that although stronger soil compactions north of Cape Cod could be slowing the northern range expansion of *Minuca pugnax*, they are clearly not stopping it. Ranges are determined by many biotic and abiotic factors (Alof et al., 2015; Burrows et al., 2014; Cassini et al., 2013; Geber, 2008; Jackson, 2009). In order to colonize new territory, they must overcome some of those barriers. We know *M. pugnax* is expanding its range (Johnson, 2014) given the populations of *M. pugnax* persisting north of Cape Cod, MA. This expansion is likely driven by the warming waters of the Gulf of Maine (GoM) (Sanford, 2006); however, their densities are lower north of Cape Cod (Johnson, 2014). Other factors, such as soil compaction, could be the reason *M. pugnax* fails to colonize northern New England marshes in the same densities as in their historical range habitat. Soil strengths between 0 and 15 psi (1.1 kg/cm^2^) (Roy et al.*, unpublished, in prep*) are more representative of saltmarshes south of the Cape, the fiddler crab’s historical range; whereas, the marshes of along the GoM can reach much greater compactions, more than 50 psi (3.51 kg/cm^2^) (Vincent, 2013).

The mechanism behind the variation between the two populations remains unclear. Our work shows that soils in marshes of the GoM are penetrable by the fiddler crabs currently found there, but in many instances might not be so for crabs from south of Cape Cod. This difference could be due to size, as Johnson et al. (2019) determined that PIE crabs are larger than southern counterparts; however, we controlled for size in our analyses and, although the largest crabs in our trials came from PIE, the majority were of comparable size (see Supplementary Results Figure 1). This suggests then that some other trait (either physically or behaviorally) influences the ability for PIE crabs to burrow in their expanded range. Our observed patterns could be due to selection on newly settled crabs coming from southern populations for only those with the ability to burrow in harder northern soils. In contrast, harder soils could lead to changes in crab phenotypes through time if these traits are plastic. Indeed, in some crab species claw morphology is even linked to water temperature (Baldridge & Smith, 2008). Both phenotypic plasticity and adaptation aid in colonization for many invasive species, which provides a compelling baseline to understand species range expansions (Smith, 2009, Stapely et al., 2015). Populations of *Minuca pugnax* in the Gulf of Maine may possess the requirements for adaptation (Sakai et al., 2001), including selective pressure (suggested by this experiment) and genetic variability (Sanford, 2006). However future studies regarding changes in population genetics should attempt to elucidate the specific mechanism driving this better burrowing capacity in PIE versus Nantucket fiddler crabs.

*M. pugnax* burrows affect the productivity, biogeochemistry, and sediment structure of their historical range saltmarshes, and so could similarly do so in ecosystems that they are expanding into (Bertness et al., 1985; Smith & Tyrrell, 2012; Johnson et al., 2020). M. pugnax can increase soil drainage, soil oxidation reduction potential, and *in situ* decomposition rates (Bertness et al., 1985, Thomas & Blum, 2010) in both the low and intermediate marsh. Degradation or loss of saltmarsh area is already exacerbated by climate change and sea level rise (Deegan et al., 2012) in PIE and other marshes north of Cape Cod. Crab burrows negatively affect the belowground growth of the soil stabilizing *S. alterniflora* (Thomas & Blum, 2010), making these low and intermediate sections of marsh the most susceptible to erosion. Bioengineers in novel habitats (such as *Minuca pugnax*) may cause additional unforeseen changes to the structure and function of PIE and other northern saltmarshes. Fortunately, our results suggest that effects in novel habitat may be predictive.

PIE crabs were capable of burrowing in harder soils, although they did not exhibit any different behavior in burrow volume once they did dig. Given that they were more likely to dig, we expected the burrowing fiddlers from PIE to displace more soil than those from Nantucket, and thus potentially affect their habitat more than Nantucket crabs. This was not the case; the burrow volume from the two populations of crabs were the same (see Figure 3). Absolute changes in soil strength due to crabs are thus likely to be similar to those in more Southern habitats. Whether the softening of harder northern marshes has comparable effects on the ecosystem relative to fiddler-induced softening of marshes south of Cape Cod remains to be seen.

Range expansions in response to global temperature increase are well documented (Davis & Shaw, 2001; Jackson et al., 2009, Loarie et al., 2009). The effect of other (sometimes subtle) physical habitat characteristics on range expanding species is not as widely explored (Brown & Vellend, 2014; Spence & Tingley, 2020). The range expansion of *Minuca pugnax* provides a wide variety of novel opportunities to study range expansions, adaptation and plasticity, and ecosystem engineering in a single species. Further, understanding if *M. pugnax* behaves as an invasive species could further clarify what impacts these range expanding crabs will have on marshes in along the Gulf of Maine through time.

## Supporting information

Supplemental Methods and Results

## Supplementary Resources

### Methods

To achieve exact compaction levels, we developed the following experimental chambers: after cutting 4, 2 cm X’s on the bottom of a clear 28 L bin (labeled E in Figure 1) for drainage, we filled each bin with 18 cm of our peat/salt water mixture (not exposed to crabs) (labeled D in Figure 1) and leveled the peat/salt throughout the bin. The weights used for compaction, were separate 28 L and 66 L bins (labeled A and B in Figure 1) filled with varying levels of sand depending on the compaction needed [no sand for 0 psi, 42.9 kg for 10 psi (0.7 kg/cm^2^), 80.8 kg for 20 psi (1.4 kg/cm^2^), and 105 kg for 25 psi (1.8 kg/cm^2^)]. The weight for the compaction process was approximated by using a formula developed by using several different weights of sand and testing what soil strength would result. A penetrometer was used to find the resulting soil strength in five places in the media, which were then averaged. The formula we developed after numerous compaction attempts to determine the weight needed is y = 4.1366x + 0.2966 (where y is the desired soil strength in psi, and x is the weight used in kg).

We cut three layers of cardboard to fit inside of the 28 L bin, which were then made into one waterproofed piece using duct tape (labeled C). The cardboard plates were placed between the peat mixture and the weight bins to evenly distribute the compression weight onto the peat/salt mixture. We nested a 16 L bin (labeled F) with 3 cm spacers (labeled G) under the 28 L bin with peat/salt to catch excess water. A black plastic bag was rolled under the water tray for easy unfurling during the experimental portion (labeled H). The weight was nested on top of the weight distribution plate and peat for 2 hours. At the end of this process, we tested whether the peat achieved the desired compaction level using a proctor style penetrometer.

**Figure 1:**
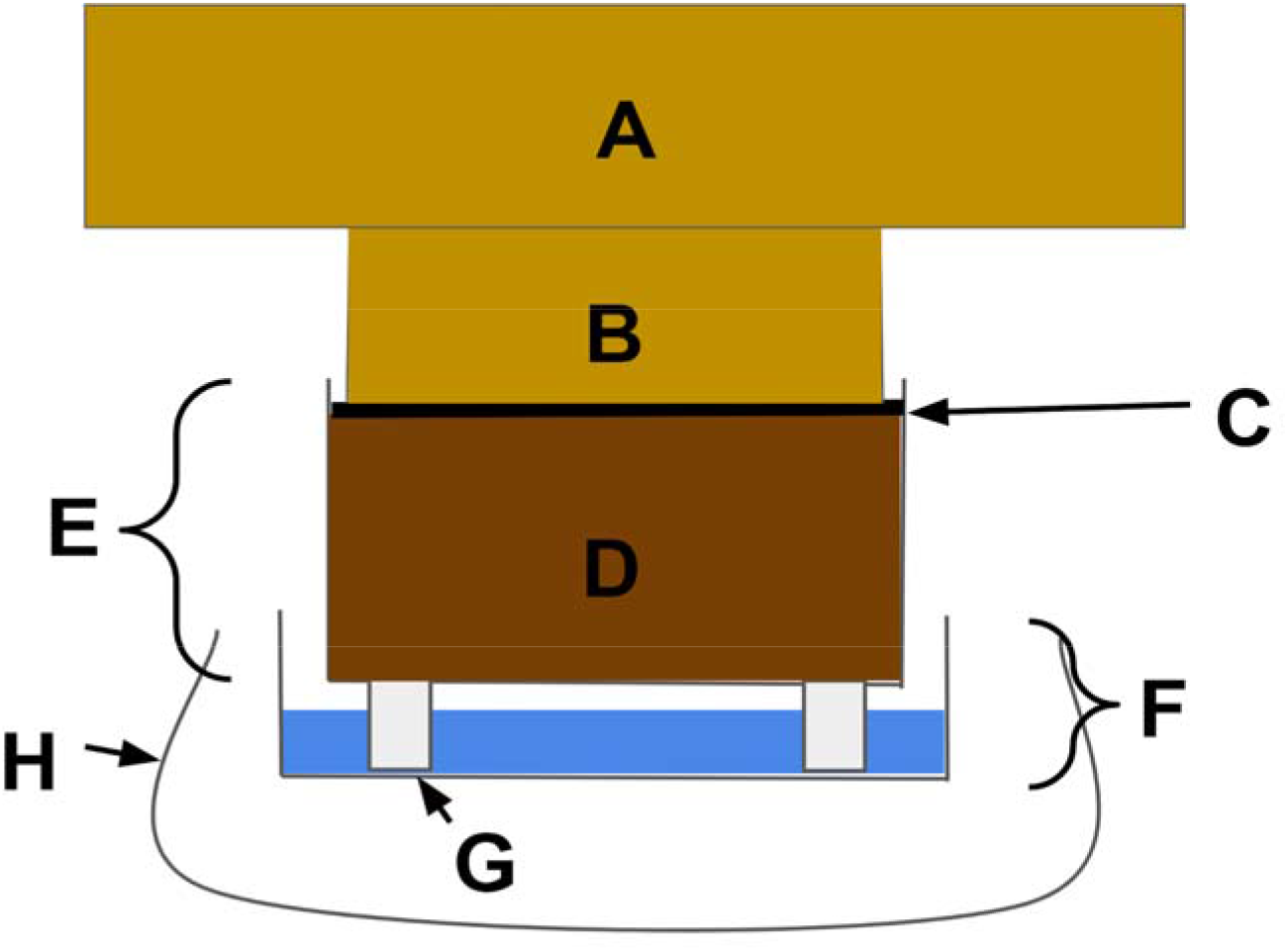
Cross section of the compression apparatus used to compact sediment. **A**: The supplementary weight, (i.e., sand), in a 66L bin that would not fit into the weight bin (B). **B**: A 28L bin filled with sand that was nested into the experimental chamber (E). **C**: A piece of cardboard with duct tape that was used to distribute the weight evenly to the peat/salt mixture. **D**: 18 cm of saltwater saturated peat moss that is being compressed. **E**: A 28L bin that is used as the experimental chamber. It has 4 drainage holes cut into the bottom to allow saltwater to escape. **F**: A 14L bin to collect excess saltwater from the experimental chamber (E). **G**: 3 cm tall plastic vials in all four corners and the center to elevate the experimental chamber and allow the media to drain. **H**: A black plastic bag to be unfurled during the experiment used to block out visual stimuli.

For the experimental portion of the trial, we attached collars to the top of each vessel (labeled I in Figure 2) to limit crab escape and minimize interference from outside the mesocosm influencing crab behavior. Each collar was made of 6 cm wide cardboard strips angled inward that were 30 cm long on the short sides of the bin, and 60 cm long on the long sides. Finally, we rolled a black plastic bag up around each vessel to further minimize outside disruption to crab behavior and mimic the light conditions in their *Spartina alterniflora* covered habitat (labeled J in Figure 2).

**Figure 2:**
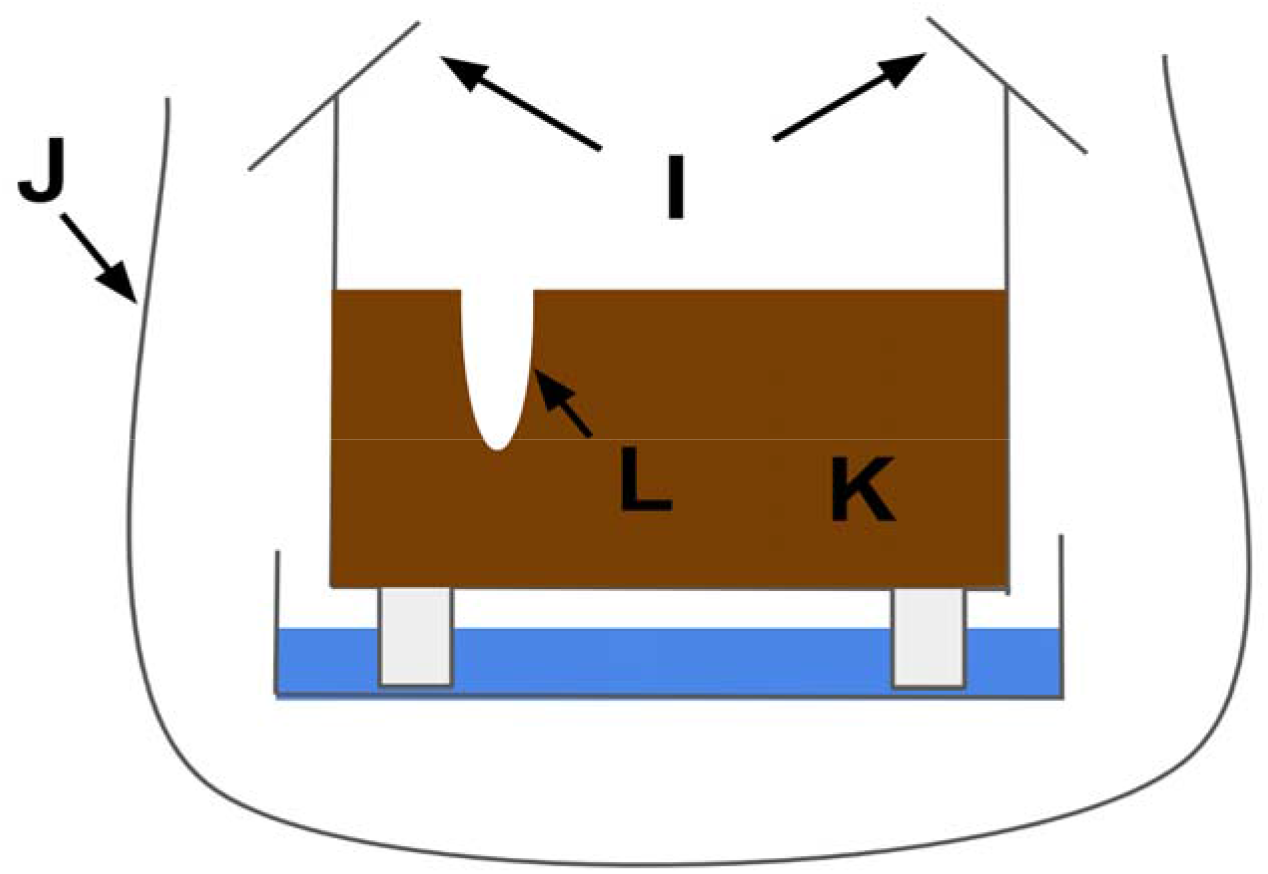
Cross section of the experimental chamber after a crab trial. **I**: Cardboard collars angled inward were used to prevent escape and minimize external stimuli. **J**: Unfurled black trash bags covered the sides of the transparent bins to further minimize stimuli and recreate light conditions under the *S. alterniflora*. **K**: Compressed peat moss. The soil strength is tested with a proctor style penetrometer before and after trials in three places in the soil. **L**: Potential crab burrows. After a trial, the crab is removed, and its mass is taken. Plaster is then poured into the burrow

**Figure 3:**
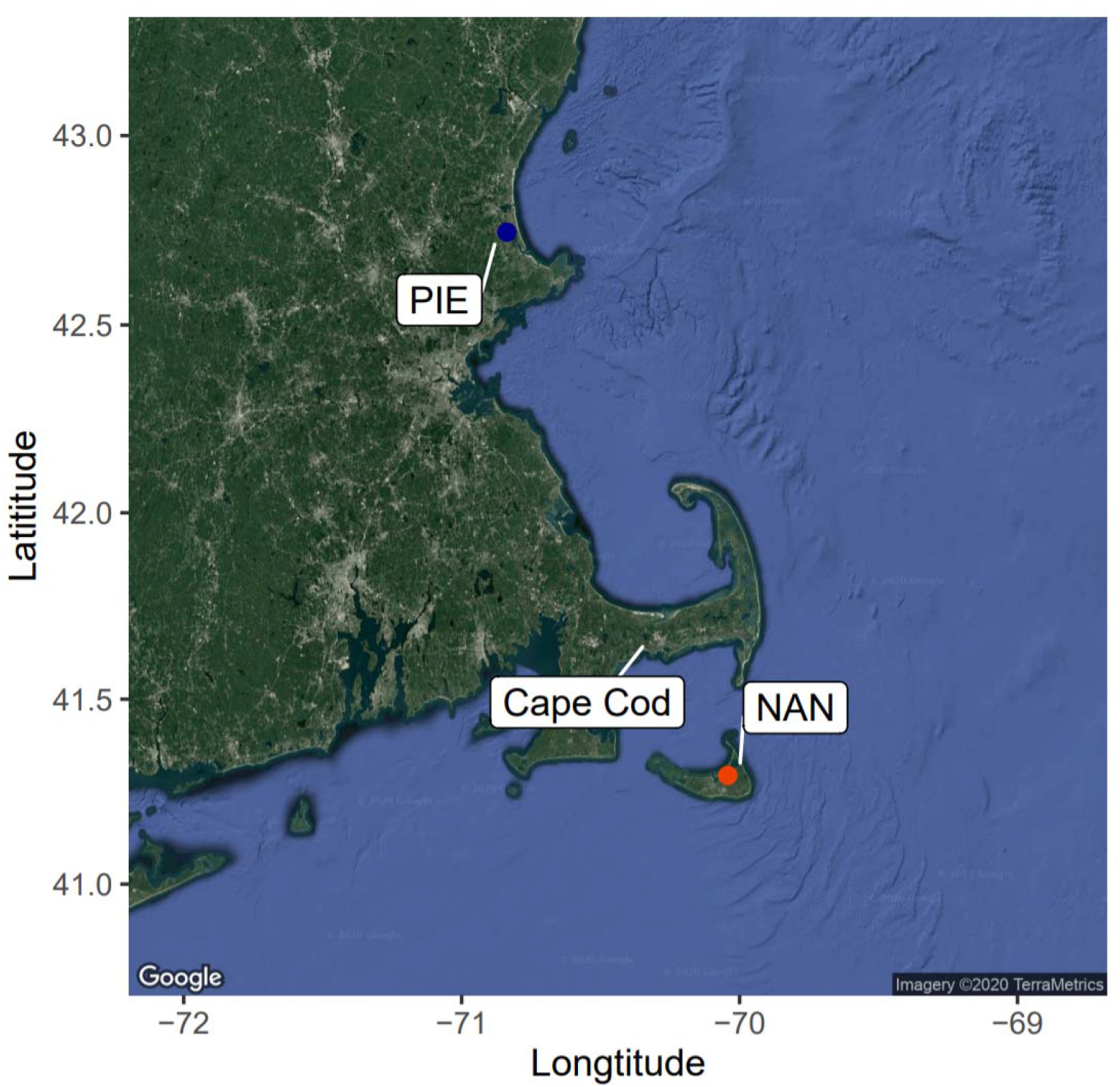
A satellite image of Massachusetts, USA showing our two collection sites. We collected at Carolton Creek in Plum Island Estuary (PIE) for fiddler crabs representing our expanded range population, and at Folger’s Marsh in Nantucket (NAN) for crabs representing their historical range. Plum Island Estuary receives relatively colder water from the north in the Gulf of Maine; Nantucket is an island in the relatively warm Gulf Stream. They are ~200 mi apart, separated by Cape Cod.

### Results

**Figure 1:**
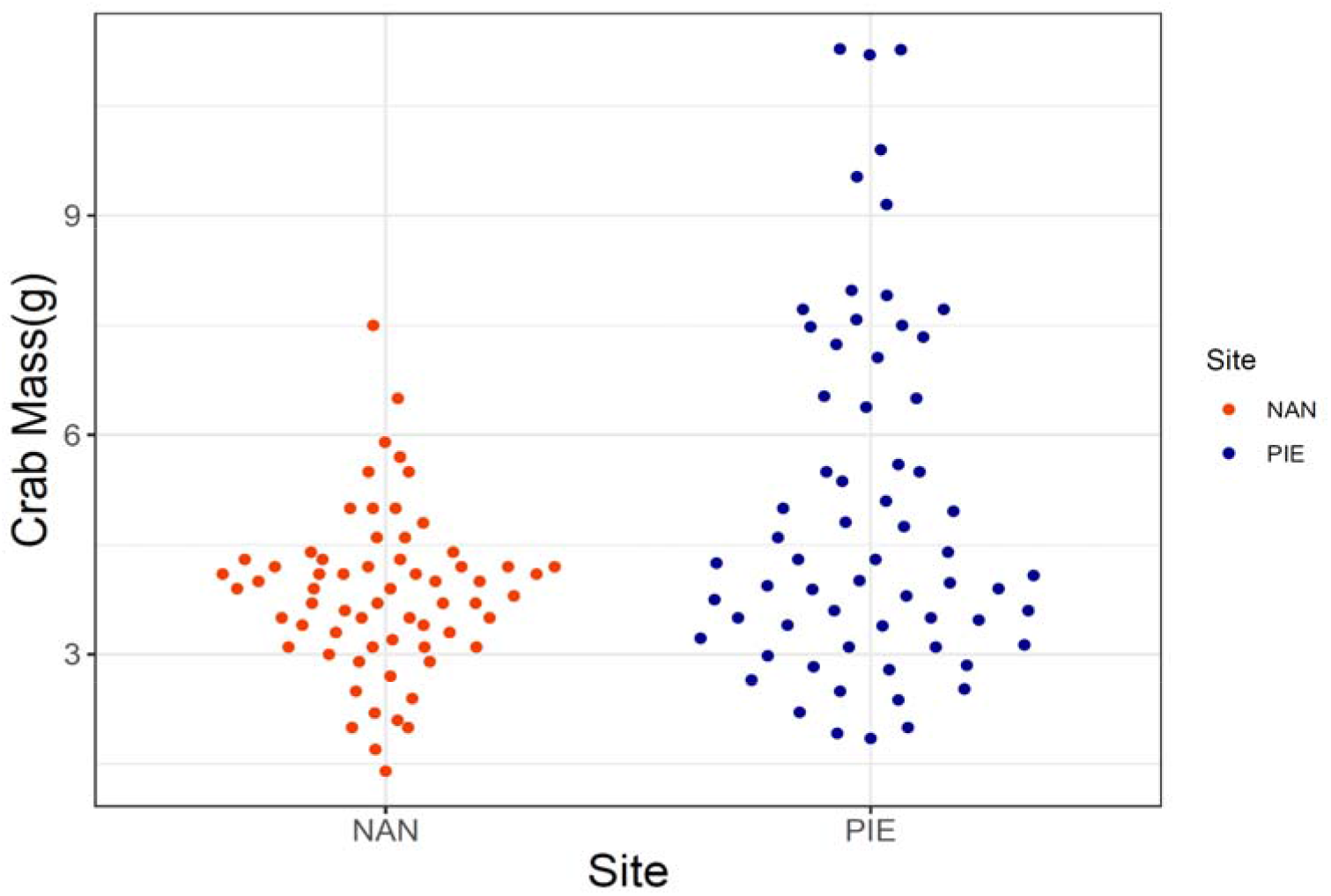
Crabs randomly collected in Plum Island Estuary were 1.22 g larger than crabs from Nantucket (p<0.001). The average mass for Nantucket crabs were 3.84 g, and Plum Island crabs were on average 5.06 g. However, the beeswarm plot above shows that most of the individuals from both Nantucket and PIE are ~3 - 4.5 g. The distribution of PIE crab individuals ranges up to 11.28 g.

## Acknowledgements

This work was supported by the National Science Foundation as part of the PIE-LTER Program (#1637630), and the Battle Fund at the University of Massachusetts – Boston Nantucket Field Station. We also would like to thank David Samuel Johnson for his insight and suggestions, as well as Abagail Kwiat and Stevens Excellent. There are no conflicts of interests to disclose.

## Notes

### Competing Interest Statement

The authors have declared no competing interest.

