## Supplemental Methods and Results for "Sediment Selection: Range Expanding Fiddler Crabs are Better Burrowers Than Their Historic Range Counterparts"

**Supplementary Resources:**

**Methods:**

To achieve exact compaction levels, we developed the following experimental chambers: after cutting 4, 2 cm X’s on the bottom of a clear 28 L bin (labeled E in Figure 1) for drainage, we filled each bin with 18 cm of our peat/salt water mixture (not exposed to crabs) (labeled D in Figure 1) and leveled the peat/salt throughout the bin. The weights used for compaction, were separate 28 L and 66 L bins (labeled A and B in Figure 1) filled with varying levels of sand depending on the compaction needed [no sand for 0 psi, 42.9 kg for 10 psi (0.7 kg/cm^2^), 80.8 kg for 20 psi (1.4 kg/cm^2^), and 105 kg for 25 psi (1.8 kg/cm^2^)]. The weight for the compaction process was approximated by using a formula developed by using several different weights of sand and testing what soil strength would result. A penetrometer was used to find the resulting soil strength in five places in the media, which were then averaged. The formula we developed after numerous compaction attempts to determine the weight needed is y = 4.1366x + 0.2966 (where **y** is the desired soil strength in psi, and **x** is the weight used in kg).

We cut three layers of cardboard to fit inside of the 28 L bin, which were then made into one waterproofed piece using duct tape (labeled C). The cardboard plates were placed between the peat mixture and the weight bins to evenly distribute the compression weight onto the peat/salt mixture. We nested a 16 L bin (labeled F) with 3 cm spacers (labeled G) under the 28 L bin with peat/salt to catch excess water. A black plastic bag was rolled under the water tray for easy unfurling during the experimental portion (labeled H). The weight was nested on top of the weight distribution plate and peat for 2 hours. At the end of this process, we tested whether the peat achieved the desired compaction level using a proctor style penetrometer.

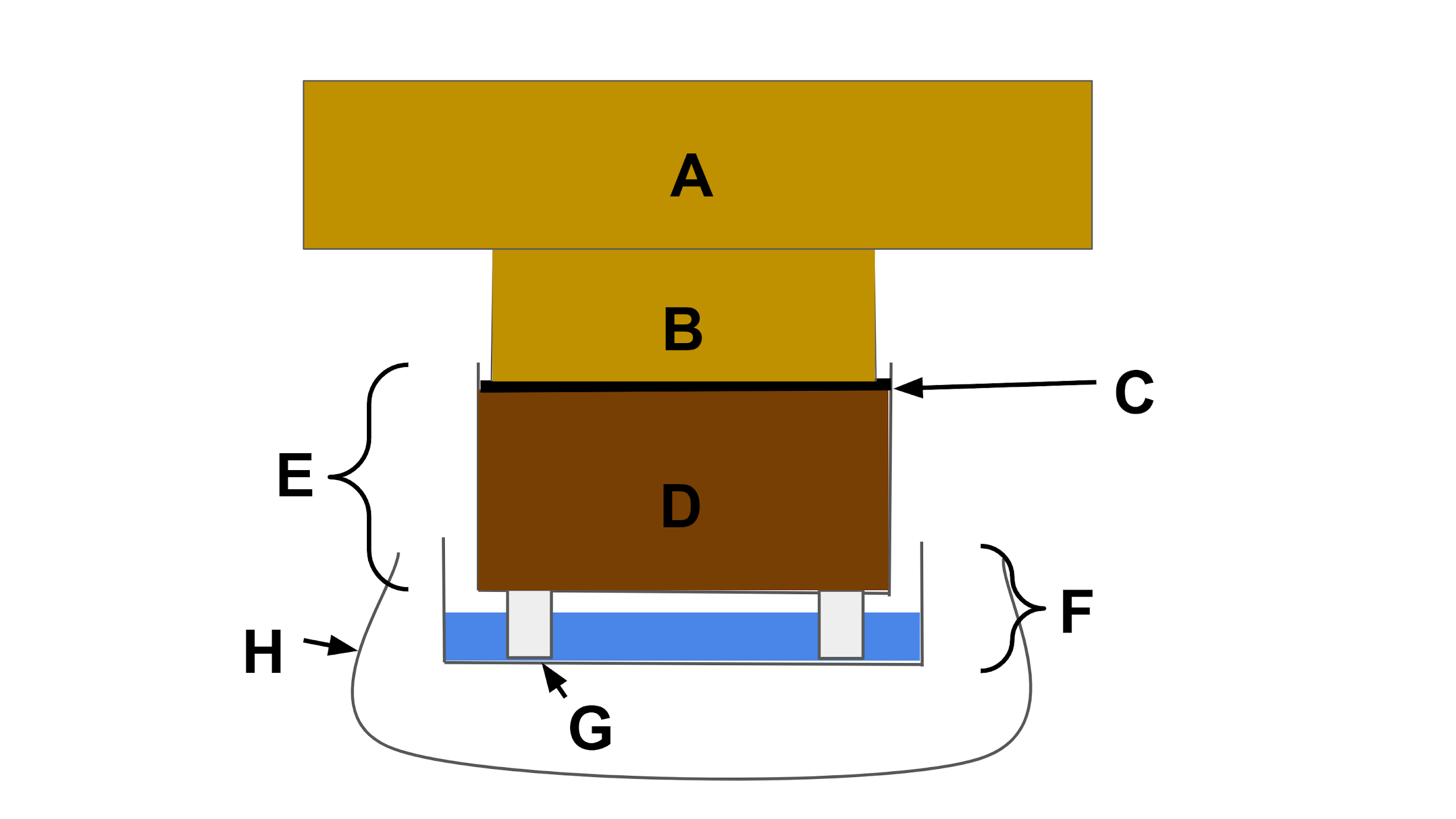

**Figure 1:** Cross section of the compression apparatus used to compact sediment. **A**: The supplementary weight, (i.e., sand), in a 66L bin that would not fit into the weight bin (B). **B**: A 28L bin filled with sand that was nested into the experimental chamber (E). **C**: A piece of cardboard with duct tape that was used to distribute the weight evenly to the peat/salt mixture. **D**: 18 cm of saltwater saturated peat moss that is being compressed. **E**: A 28L bin that is used as the experimental chamber. It has 4 drainage holes cut into the bottom to allow saltwater to escape. **F**: A 14L bin to collect excess saltwater from the experimental chamber (E). **G**: 3 cm tall plastic vials in all four corners and the center to elevate the experimental chamber and allow the media to drain. **H**: A black plastic bag to be unfurled during the experiment used to block out visual stimuli.

For the experimental portion of the trial, we attached collars to the top of each vessel (labeled I in Figure 2) to limit crab escape and minimize interference from outside the mesocosm influencing crab behavior. Each collar was made of 6 cm wide cardboard strips angled inward that were 30 cm long on the short sides of the bin, and 60 cm long on the long sides. Finally, we rolled a black plastic bag up around each vessel to further minimize outside disruption to crab behavior and mimic the light conditions in their *Spartina alterniflora* covered habitat (labeled J in Figure 2).

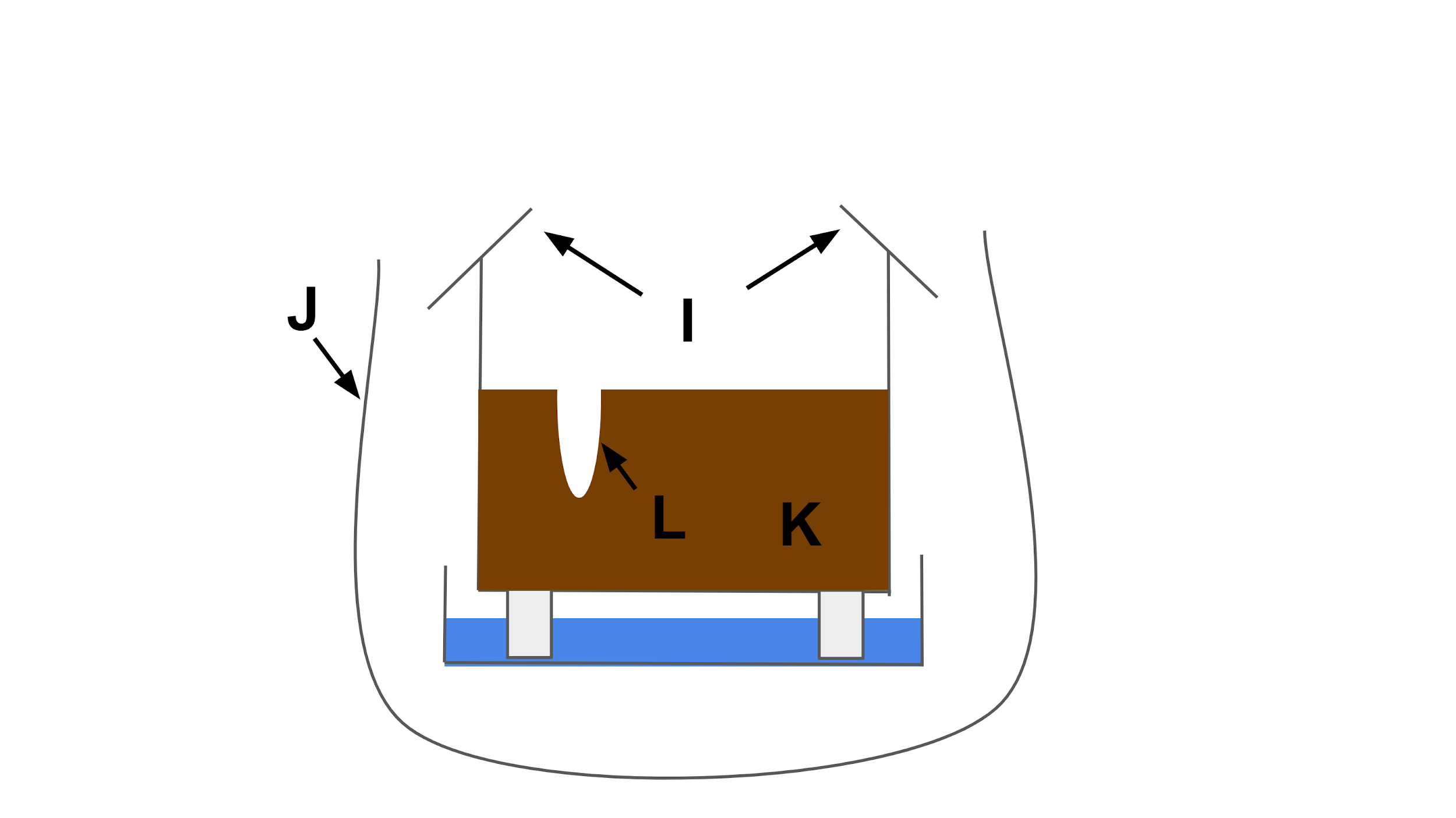

**Figure 2:** Cross section of the experimental chamber after a crab trial. **I**: Cardboard collars angled inward were used to prevent escape and minimize external stimuli. **J**: Unfurled black trash bags covered the sides of the transparent bins to further minimize stimuli and recreate light conditions under the *S. alterniflora.* **K**: Compressed peat moss. The soil strength is tested with a proctor style penetrometer before and after trials in three places in the soil. **L**: Potential crab burrows. After a trial, the crab is removed, and its mass is taken. Plaster is then poured into the burrow

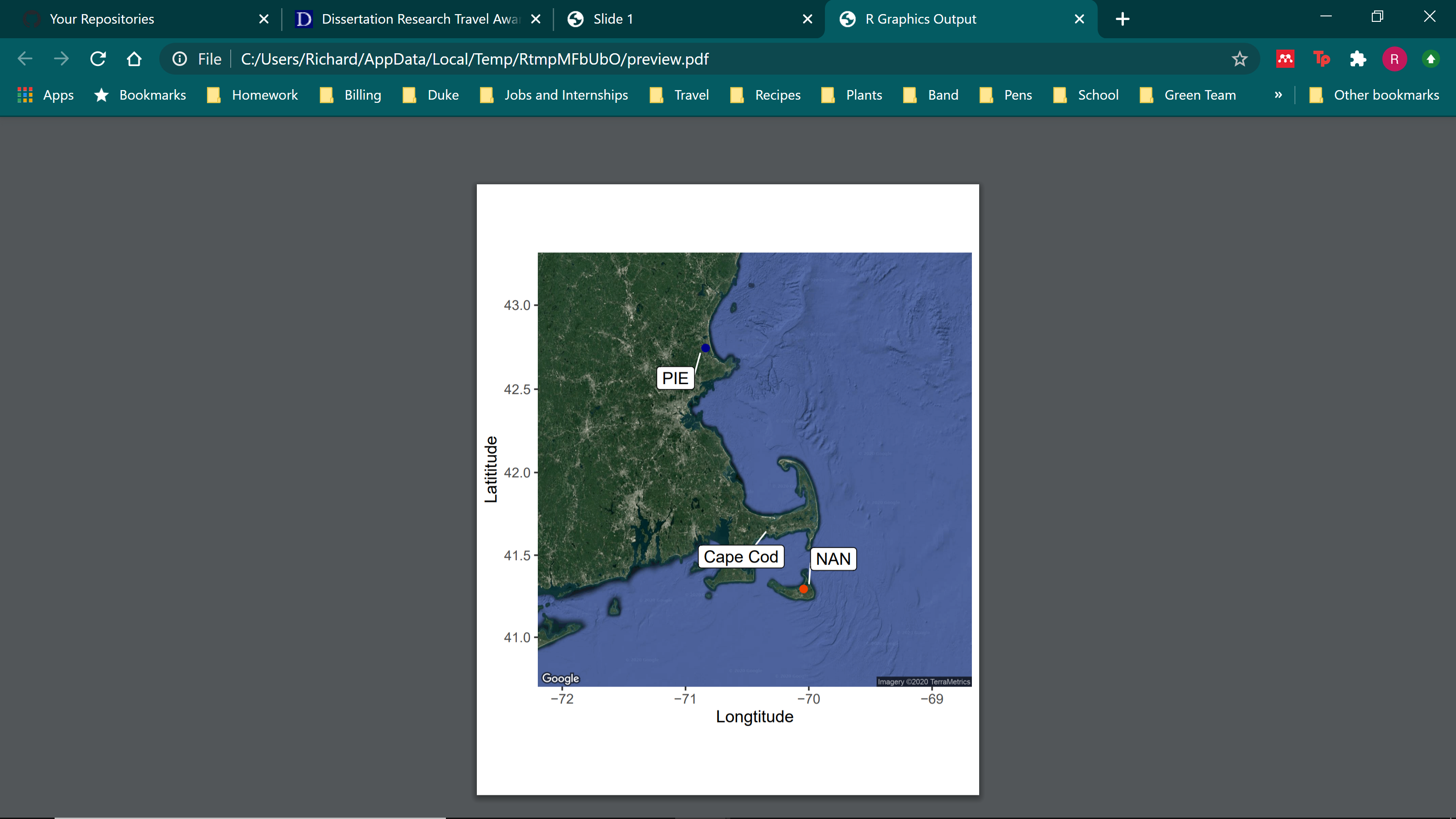

**Figure 3:** A satellite image of Massachusetts, USA showing our two collection sites. We collected at Carolton Creek in Plum Island Estuary (PIE) for fiddler crabs representing our expanded range population, and at Folger’s Marsh in Nantucket (NAN) for crabs representing their native range. Plum Island Estuary receives relatively colder water from the north in the Gulf of Maine; Nantucket is an island in the relatively warm Gulf Stream. They are ~200 mi apart, separated by Cape Cod.

**Results:**

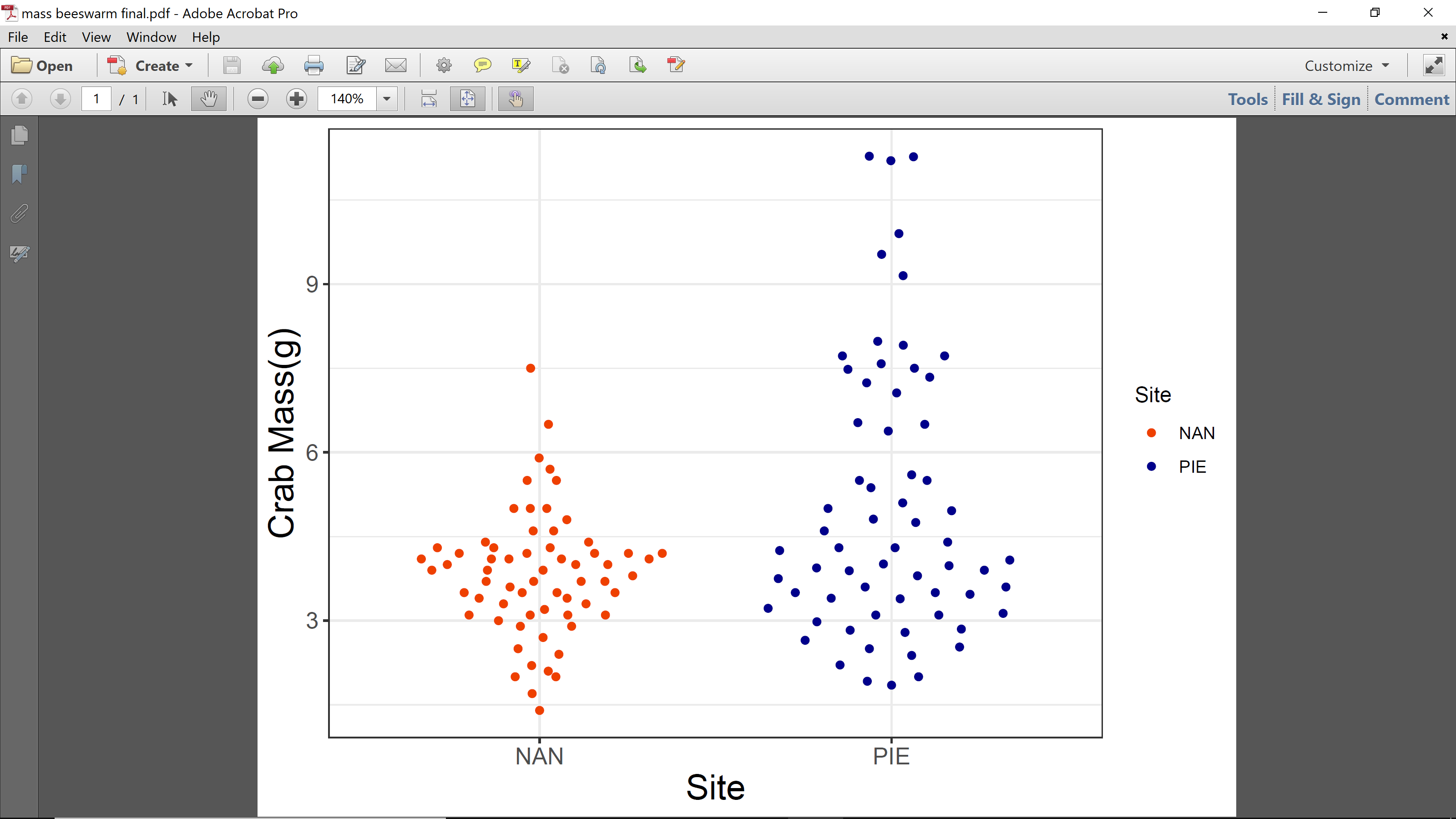

**Figure 1:** Crabs randomly collected in Plum Island Estuary were 1.22 g larger than crabs from Nantucket (p<0.001). The average mass for Nantucket crabs were 3.84 g, and Plum Island crabs were on average 5.06 g. However, the beeswarm plot above shows that most of the individuals from both Nantucket and PIE are ~3 - 4.5 g. The distribution of PIE crab individuals ranges up to 11.28 g.
